# Changes of urinary proteome in rats after intragastric administration of zinc gluconate

**DOI:** 10.1101/2024.03.04.583149

**Authors:** Ziyun Shen, Minhui Yang, Haitong Wang, Youhe Gao

**Affiliations:** Gene Engineering Drug and Biotechnology Beijing Key Laboratory, College of Life Sciences, Beijing Normal University, Beijing 100875, China

**Keywords:** zinc, urine, proteome, zinc gluconate, nutrients, mineral elements

## Abstract

Zinc is an essential element for maintaining normal physiological function in living organisms. In this study, the urine proteome of rats before and after short-term intragastric administration of 82 mg/kg/d zinc gluconate (equivalent to 11.7 mg/kg/d zinc) was compared and analyzed. Many differential proteins have been reported to be zinc-related, such as mucin-2 (MUC-2) (14 times before compared with after gavage, p = 0.005) and transthyretin (3.9 times after gavage compared with before gavage, p = 0.0004). Biological processes enriched in differential proteins (e.g., regulation of apoptosis process, immune system process, etc.), molecular functions (e.g., calcium binding, copper binding, signaling receptor activity, etc.), KEGG pathways (e.g., complement and coagulation cascades, PI3K-Akt signaling pathway, etc.) showed correlation with zinc. In this study, we explore the overall effect of zinc on the body from the perspective of urine proteomics, which is helpful to deeply understand the biological function of zinc and broaden the application potential of urine proteomics.

## 1 Introduction

Zinc plays a crucial role in various physiological processes. Zinc is a structural component and a catalytic and regulatory cofactor of many enzymes and transcription factors, and is involved in DNA synthesis, protein synthesis, proliferation, maturation, death, immune response, antioxidant defense, and other life processes in cells^[1,2]^. Zinc also acts as a regulatory and signal transduction element in intercellular and intracellular communication^[3]^.

Zinc deficiency can lead to a variety of health problems, including growth retardation, immunodeficiency, hypogonadism, and neuronal and sensory dysfunctions^[4]^. Abnormalities in zinc homeostasis have also been linked to the development of chronic diseases such as cancer, diabetes, depression, Wilson’s disease, and Alzheimer’s disease^[5]^. Zinc homeostasis is regulated by zinc transporter proteins and metallothioneins.

Since urine is not part of the internal environment, in contrast to plasma, there is no mechanism for homeostasis, and it is able to accumulate early changes in the physiological state of the organism, reflecting more sensitively the changes in the organism, and is a source of next-generation biomarkers^[6]^. The proteins in urine contain a wealth of information that can reflect small changes produced in different systems and organs of the organism. Our laboratory has previously reported that the urine proteome is able to reflect the effects of magnesium threonate intake on the body in a more systematic and comprehensive manner, and has the potential to provide clues for clinical nutrition research and practice^[7]^.

Although the physiological functions of zinc have been extensively studied, to date, there have been no studies exploring the overall effect of zinc on the organism from the perspective of the urinary proteome. In this study, zinc gluconate was chosen as the supplement for the study population. Zinc gluconate is an organic zinc supplement with low gastric mucosal irritation, easy absorption in the body, and high absorption and solubility, which is widely used in health care products, pharmaceuticals and food.

The aim of this study was to investigate the changes in the urinary proteome of rats after zinc gluconate intake, in the hope of deepening the understanding of the physiological functions of zinc, providing new perspectives and new clues for nutritional studies, and contributing to a more scientific guidance for human health and dietary regulation of trace elements.

## 2 Materials and Methods

### 2.1 Experimental materials

#### 2.1.1 Experimental consumables

5 ml sterile syringe (BD), gavage needle (16-gauge, 80 mm, curved needle), 1.5 ml/2 ml centrifuge tube (Axygen, USA), 50 ml/15 ml centrifuge tube (Corning, USA), 96-well cell culture plate (Corning, USA), 10 kD filter (Pall, USA), Oasis HLB solid phase extraction column (Waters, USA), 1ml/200ul/20ul pipette tips (Axygen, USA), BCA kit (Thermo Fisher Scientific, USA), high pH reverse peptide separation kit (Thermo Fisher Scientific, USA), iRT ( indexed retention time, BioGnosis, UK).

#### 2.1.2 Experimental apparatus

Rat metabolic cages (Beijing Jiayuan Xingye Science and Technology Co., Ltd.), frozen high-speed centrifuge (Thermo Fisher Scientific, USA), vacuum concentrator (Thermo Fisher Scientific, USA), DK-S22 electric thermostatic water bath (Shanghai Jinghong Experimental Equipment Co., Ltd.), full-wavelength multifunctional enzyme labeling instrument (BMG Labtech, Germany), oscillator (Thermo Fisher Scientific, USA), TS100 constant temperature mixer (Hangzhou Ruicheng Instruments Co. BMG Labtech), electronic balance (METTLER TOLEDO, Switzerland), -80 □ ultralow-temperature freezing refrigerator (Thermo Fisher Scientific, USA), EASY-nLC1200 Ultra High Performance Liquid Chromatography system (Thermo Fisher Scientific, USA), and Orbitrap Fusion Lumos Tribird Mass Spectrometer (Thermo Fisher Scientific, USA) were used.

#### 2.1.3 Experimental reagents

Zinc gluconate (Zinc gluconate) was purchased from Shanghai Yuanye Biotechnology Co., Ltd, CAS No. 4468-02-4, molecular formula C12H22ZnO14, purity over 98%. In addition, Trypsin Golden (Promega, USA), dithiothreitol (DTT) (Sigma, Germany), iodoacetamide (IAA) (Sigma, Germany), ammonium bicarbonate NH4HCO3 (Sigma, Germany), urea (Sigma, Germany), and purified water (Wahaha, China) were used, Methanol for mass spectrometry (Thermo Fisher Scientific, USA), Acetonitrile for mass spectrometry (Thermo Fisher Scientific, USA), Purified water for mass spectrometry (Thermo Fisher Scientific, USA), Tris-Base (Promega, USA), Thiourea (Sigma, Germany) and other reagents.

#### 2.1.4 Analysis software

Proteome Discoverer (Version2.1, Thermo Fisher Scientific, USA), Spectronaut Pulsar (Biognosys, UK), Ingenuity Pathway Analysis (Qiagen, Germany); R studio (Version1.2.5001); Xftp 7; Xshell 7.

### 2.2 Experimental Methods

#### 2.2.1 Animal modeling

The present study was conducted using 17-week-old rats to minimize the effects of growth and development during gavage. Five healthy SD (Sprague Dawley) 9-week-old male rats (250±20 g) were purchased from Beijing Viton Lihua Laboratory Animal Technology Co. The rats were kept in a standard environment (room temperature (22±2)□, humidity 65%-70%) for 8 weeks and weighed 500-600 g. The experiments were started, and all experimental operations followed the review and approval of the Ethics Committee of the School of Life Sciences, Beijing Normal University.

Dietary nutrient tolerable upper intake levels (UL, tolerable upper intake levels): the average daily maximum intake of a nutrient for a population of a certain physiological stage and gender, which has no side effects or risks to the health of almost all individuals. Recommended nutrient intakes (RNI): intake levels that meet 97-98% of the individual needs of a particular age, gender, or physiological group.

According to the Dietary Guidelines for Chinese Residents, the UL for zinc is 40 mg/d^[8]^. The UL for humans was converted to a dose for rats based on body surface area and body weight equal to approximately 3.6 mg/kg-d, i.e., 25.3 mg/kg-d of zinc gluconate. In the present study, the dose of zinc in rats was 11.7 mg/kg-d by gavage, and that of zinc gluconate was 82 mg/kg-d, which was three times the UL. 4.2 g of zinc gluconate was dissolved in 500 ml of sterile water and configured as a gavage solution. After one week of normal feeding, each rat was gavaged with 5 ml of zinc gluconate solution once a day for 4 consecutive days to establish a high zinc model. The first day of gavage was recorded as Zn-D1, and so on. Sampling time points were set before and after gavage for their own before and after control. The samples collected on the day before gavage were the control group, recorded as Zn-D0, and the samples were numbered 36-40, and the samples collected on the 4th day of gavage were the experimental group, recorded as Zn-D4, and the samples were numbered 46-50.

**Fig. 1.**
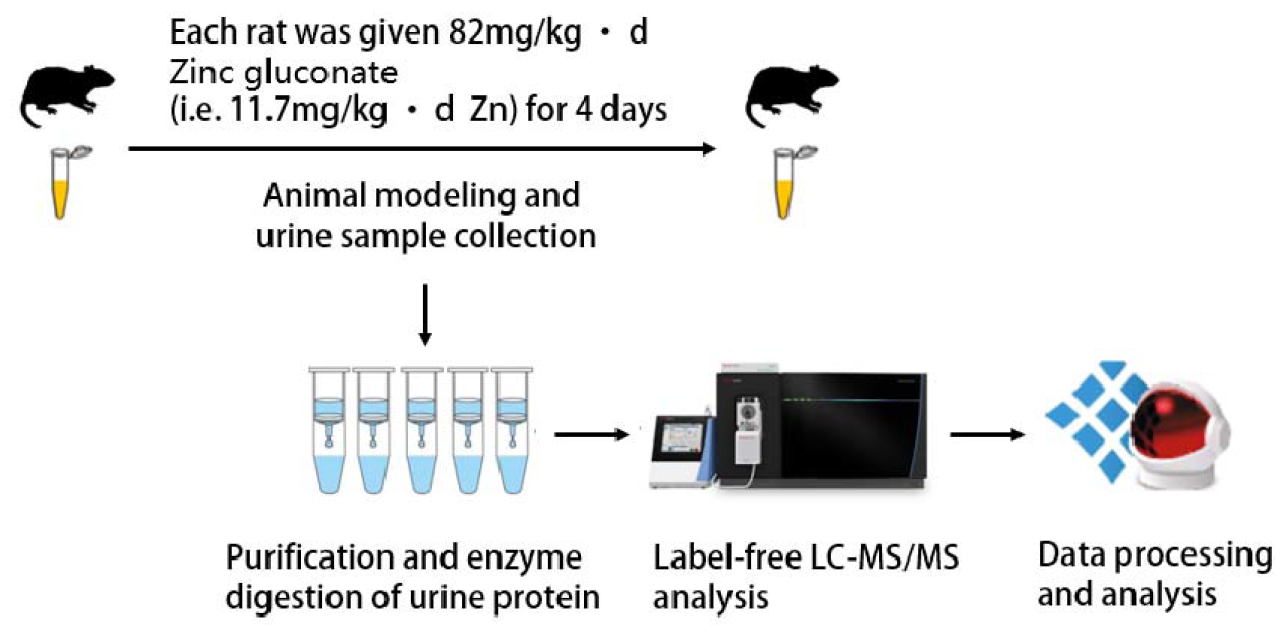
Research methodology and technical route.

#### 2.2.2 Urine sample collection

One day before the start of gavage mineral supplementation (D0) and 4 days after gavage mineral supplementation (D4), each rat was individually placed in a metabolic cage at the same time, fasted and dehydrated for 12 h. Urine was collected overnight, and the urine samples were collected and placed in the refrigerator at -80°C for temporary storage.

#### 2.2.3 Urine sample processing

Two milliliters of urine sample was removed, thawed, and centrifuged at 4 °C and 12,000×g for 30 minutes. The supernatant was removed, and 1 M dithiothreitol (DTT, Sigma) storage solution (40 µl) was added to reach the working concentration of DTT (20 mM). The solution was mixed well and then heated in a metal bath at 37 °C for 60 minutes and allowed to cool to room temperature.

Then, iodoacetamide (IAA, Sigma) storage solution (100 µl) was added to reach the working concentration of IAM, mixed well and then reacted for 45 minutes at room temperature protected from light. At the end of the reaction, the samples were transferred to new centrifuge tubes, mixed thoroughly with three times the volume of precooled anhydrous ethanol, and placed in a freezer at -20 °C for 24 h to precipitate the proteins.

At the end of precipitation, the sample was centrifuged at 4 °C for 30 minutes at 10,000×g, the supernatant was discarded, the protein precipitate was dried, and 200 µl of 20 mM Tris solution was added to the protein precipitate to reconstitute it. After centrifugation, the supernatant was retained, and the protein concentration was determined by the Bradford method.

Using the filter-assisted sample preparation (FASP) method, urinary protein extracts were added to the filter membrane of a 10-kD ultrafiltration tube (Pall, Port Washington, NY, USA) and washed three times with 20 mM Tris solution. The protein was resolubilized by the addition of 30 mM Tris solution, and the protein was added in a proportional manner (urinary protein:trypsin = 50:1) to each sample. Trypsin (Trypsin Gold, Mass Spec Grade, Promega, Fitchburg, WI, USA) was used to digest proteins at 37 °C for 16 h.

The digested filtrate was the peptide mixture. The collected peptide mixture was desalted by an Oasis HLB solid phase extraction column, dried under vacuum, and stored at -80 °C. The peptide mixture was then extracted with a 0.1% peptide mixture. The lyophilized peptide powder was redissolved by adding 30 μL of 0.1% formic acid water, and then the peptide concentration was determined by using the BCA kit. The peptide concentration was diluted to 0.5 μg/μL, and 4 μL of each sample was removed as the mixed sample.

#### 2.2.4 LC-MS/MS analysis

All identification samples were added to a 100-fold dilution of iRT standard solution at a ratio of 20:1 sample:iRT, and the retention times were standardized. Data-independent acquisition (DIA) was performed on all samples, and each sample measurement was repeated 3 times, with 1-mix samples inserted after every 10 runs as a quality control. The 1-µg samples were separated using EASY-nLC1200 liquid chromatography (elution time: 90 min, gradient: mobile phase A: 0.1% formic acid, mobile phase B: 80% acetonitrile), the eluted peptides were entered into the Orbitrap Fusion Lumos Tribird mass spectrometer for analysis, and the corresponding raw files of the samples were generated.2.2.5 Data processing and analysis

The raw files collected in DIA mode were imported into Spectronaut software for analysis, and the highly reliable protein standard was peptide q value<0.01. The peak area quantification method was applied to quantify the protein by applying the peak area of all fragmented ion peaks of secondary peptides, and the automatic normalization was processed.

Proteins containing two or more specific peptides were retained, and missing values were replaced with 0. The amount of different proteins identified in each sample was calculated, and samples from rats before gavage of mineral supplements were compared with samples from rats 4 days after gavage of mineral supplements to screen for differential proteins.

Unsupervised cluster analysis (HCA), principal component analysis (PCA), and OPLS-DA analysis were performed using the Wukong platform (https://omicsolution.org/wkomics/main/). Functional enrichment analysis of differential proteins was performed using the DAVID database (https://david.ncifcrf.gov/) to obtain results in 3 areas: biological process, cellular localization and molecular function. Differential proteins and related pathways were searched based on Pubmed database (https://pubmed.ncbi.nlm.nih.gov/). Protein interaction network analysis was performed using the STRING database (https://cn.string-db.org/).

## 3. Results and Discussion

### 3.1 Differential protein analysis

The missing values were replaced with 0. 112 differential proteins were screened by comparing the pre-gavage samples of rats with the samples on day 4 of gavage. Differential proteins were screened under the following conditions: p-value < 0.05 by t-test analysis, and Fold change (FC) > 1.5 or < 0.67. Sequence numbers of 5 differential proteins were not queried in Uniprot, probably because the entries had been deleted or merged.

Using the Uniprot database, Protein Accessions was entered into the search box to download the name function, GO analysis results of the proteins. Using PubMed database to analyze the protein function of differential proteins and literature search, the relationship between differential proteins and zinc was analyzed in detail one by one. The specific method is as follows: enter the protein name of the differential protein together with zinc into the Pubmed search box, and the search range is title/abstract, for example, “zinc [Title/Abstract] AND Protein [Title/Abstract]”. The literature was then read to confirm the relationship between the differential proteins and zinc. The 107 differential proteins and related literature are listed in Table 1.

**Table 1.**
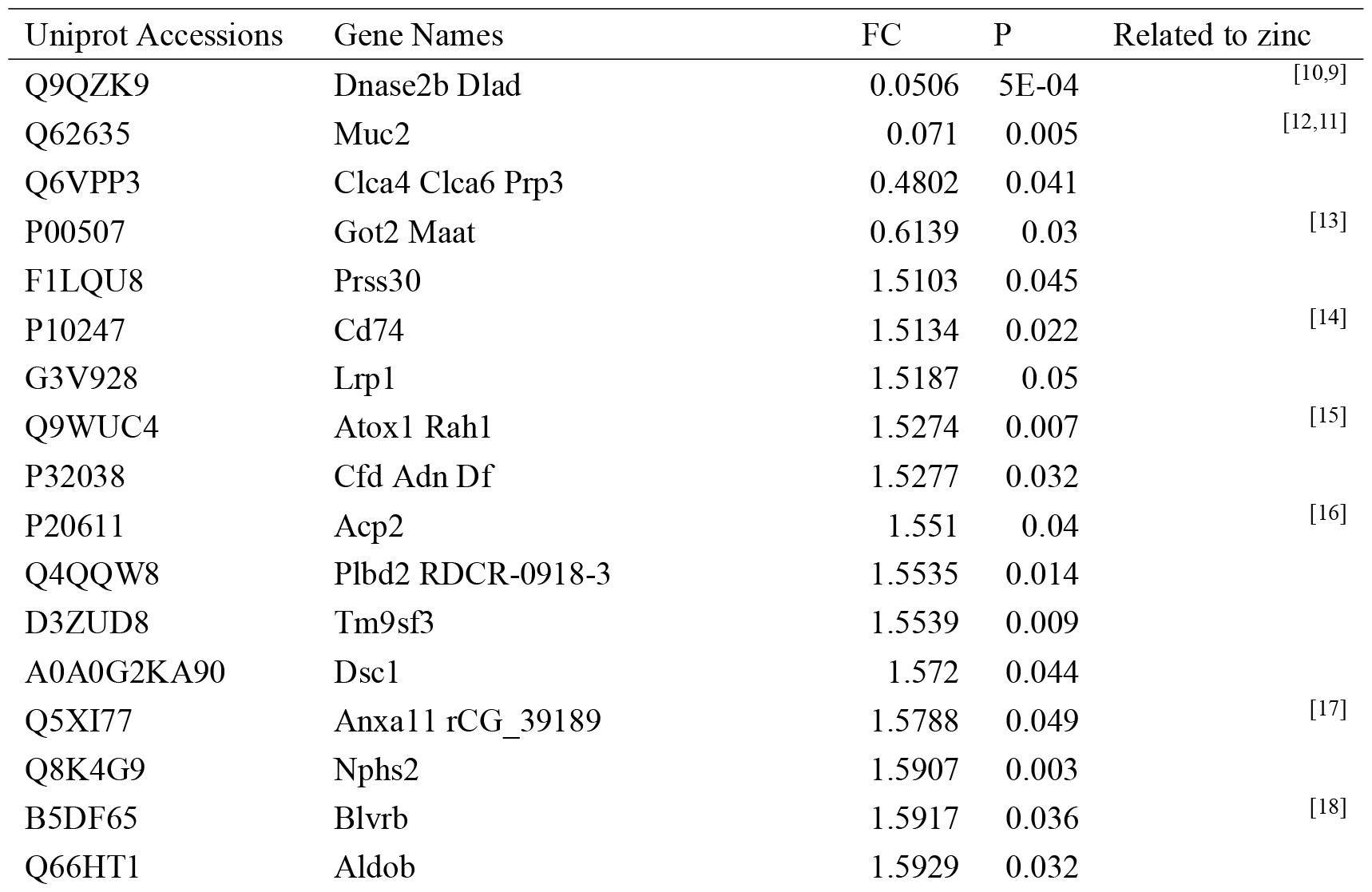
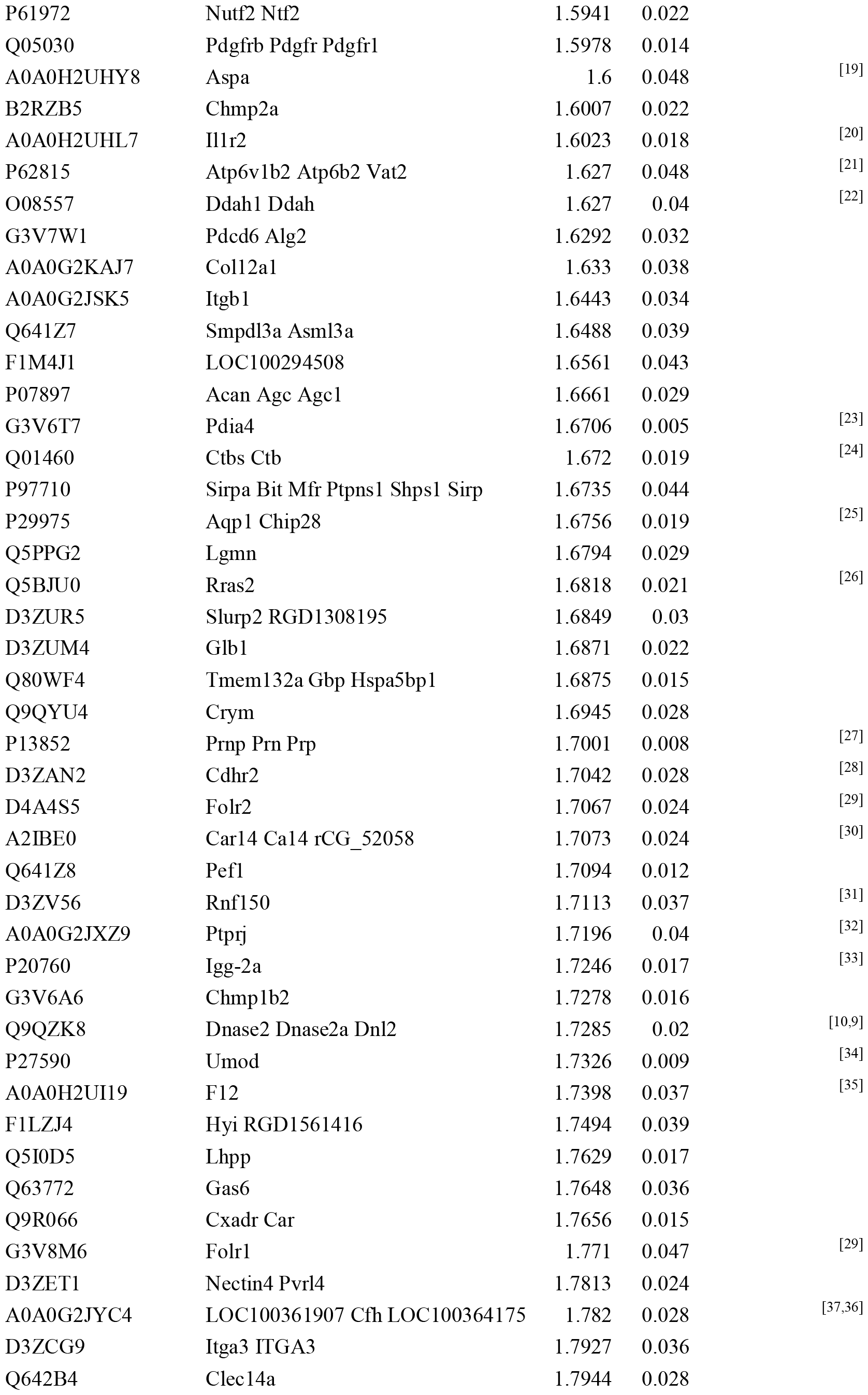
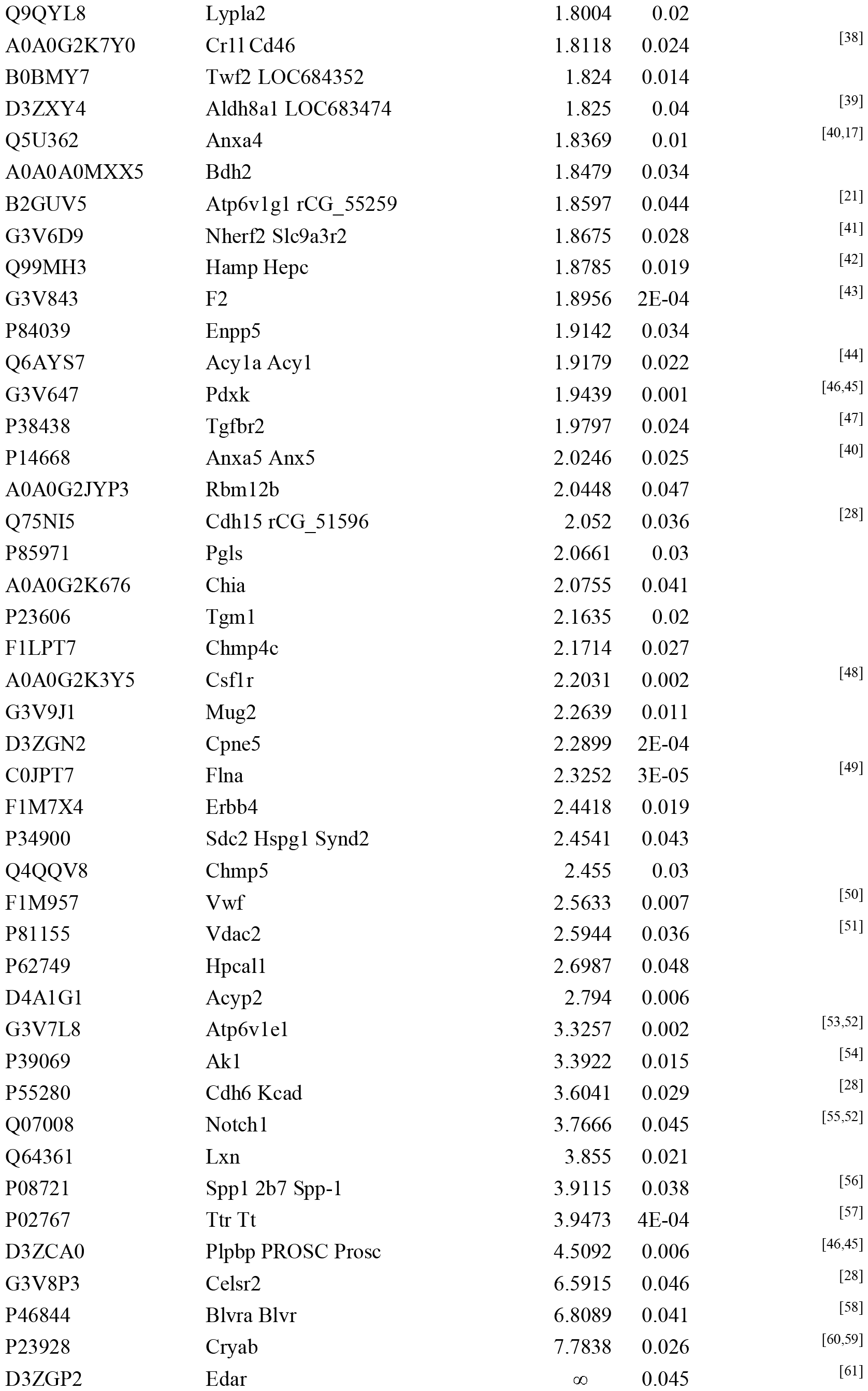
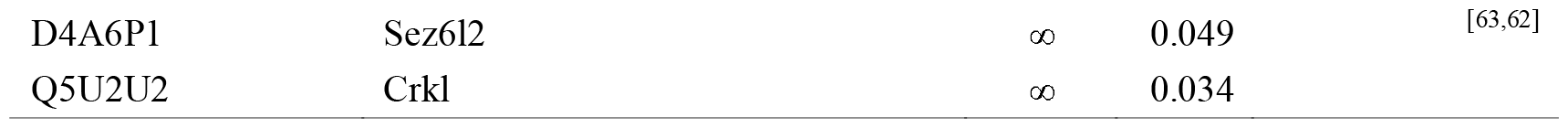
Differential proteins in the comparative analysis of the Zn-D0 and Zn-D4 groups (p-value < 0.05, FC > 1.5 or < 0.67)

According to Uniprot analysis, many differential proteins have the function of binding zinc ions, including acid sphingomyelinase-like phosphodiesterase 3a, membrane-bound carbonic anhydrase 14, ectonucleotide pyrophosphatase/phosphodiesterase family member 5 (E-NPP), and Biliverdin reductase A (BVR A). Hepcidin (FC=1.9, p=0.019) are involved in biological processes including the response to zinc ions. Zinc may be involved in the regulation of hepcidin production^[42]^ .

Zinc is a structural component of many proteins (e.g., a variety of enzymes and transcription factors) and is a regulatory cofactor for a number of differential proteins that modulate protein activity. For example, dimethylarginine dimethylaminohydrolase 1 (DDAH-1) is inhibited by zinc ions. DDAH-1 is downregulated in the hippocampus of zinc-deficient rats^[22]^. The

There were 20 differential proteins that met p<0.01, most of which were reported to be associated with zinc.

Deoxyribonuclease-2-β had an FC of 0.05 and a p-value of 0.0005. Deoxyribonuclease II is activated by intracellular acidification that occurs during apoptosis, and zinc inhibits intracellular acidification associated with apoptosis, thereby inhibiting deoxyribonuclease II^[10]^. Mucin-2 (MUC-2) had an FC of 0.07 and a p-value of 0.005. Increased abundance of MUC2 was found in the jejunum of the offspring of mothers on a high-zinc diet^[12]^.

The major prion protein (PrP) (FC=1.7,p=0.008) may be involved in neuronal zinc homeostasis^[27]^. There is a correlation between transthyretin (FC=3.9,p=0.0004) and zinc^[57]^.

Pyridoxal phosphate homeostasis protein (PPSP) has an FC of 4.5 and a p-value of 0.006. pyridoxal kinase (PK) has an FC of 1.9 and a p-value of 0.001. At physiological concentrations, zinc stimulates pyridoxal kinase activity and promotes pyridoxal phosphate formation^[46]^.

The FC for three differential proteins, Seizure related 6 homolog like 2, Ectodysplasin-A receptor, and Crk-like protein, was ∞, i.e., this protein was not detected in the pre-gastric sample and was detected in the post-gastric sample. protein was not detected in the pre-gastric samples, while it was detected in the post-gastric samples. According to the literature, zinc is associated with neuronal damage and death after seizures^[62,63]^. The gene edar is associated with inflammation and specifically responds to zinc oxide nanoparticles (ZnO NPs)^[61]^. Due to space limitations, only a few examples are listed, and the differential proteins, as well as the relevant literature, are detailed in Table 1.

### 3.2 Biological pathway analysis

Gene Oncology (GO) analysis of 112 differential proteins (P-value < 0.05, FC > 1.5 or < 0.67) using the DAVID database enriched to 83 biological processes (BPs) (P-value < 0.05), as shown in Table 2.

**Table 2.**
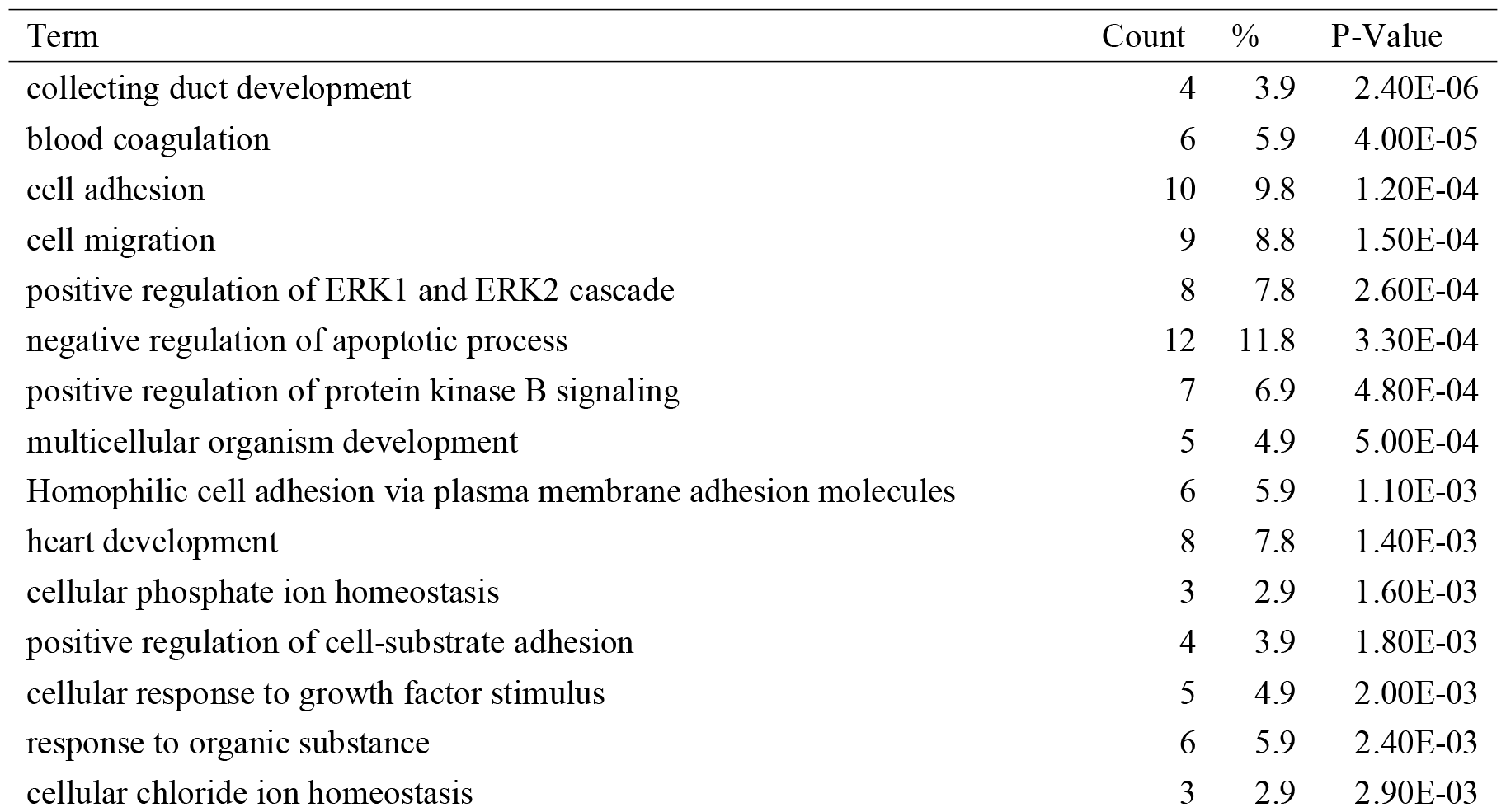
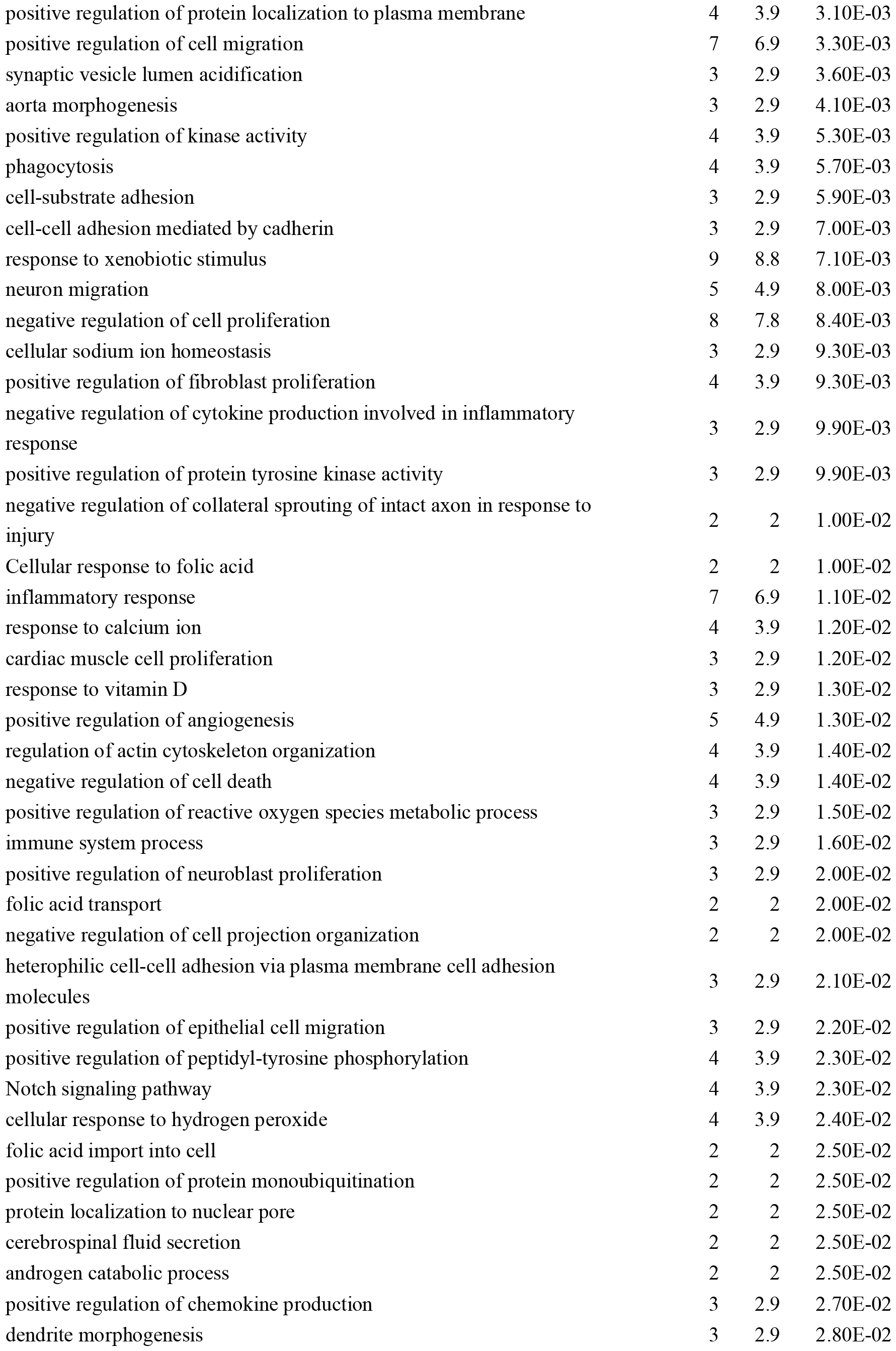
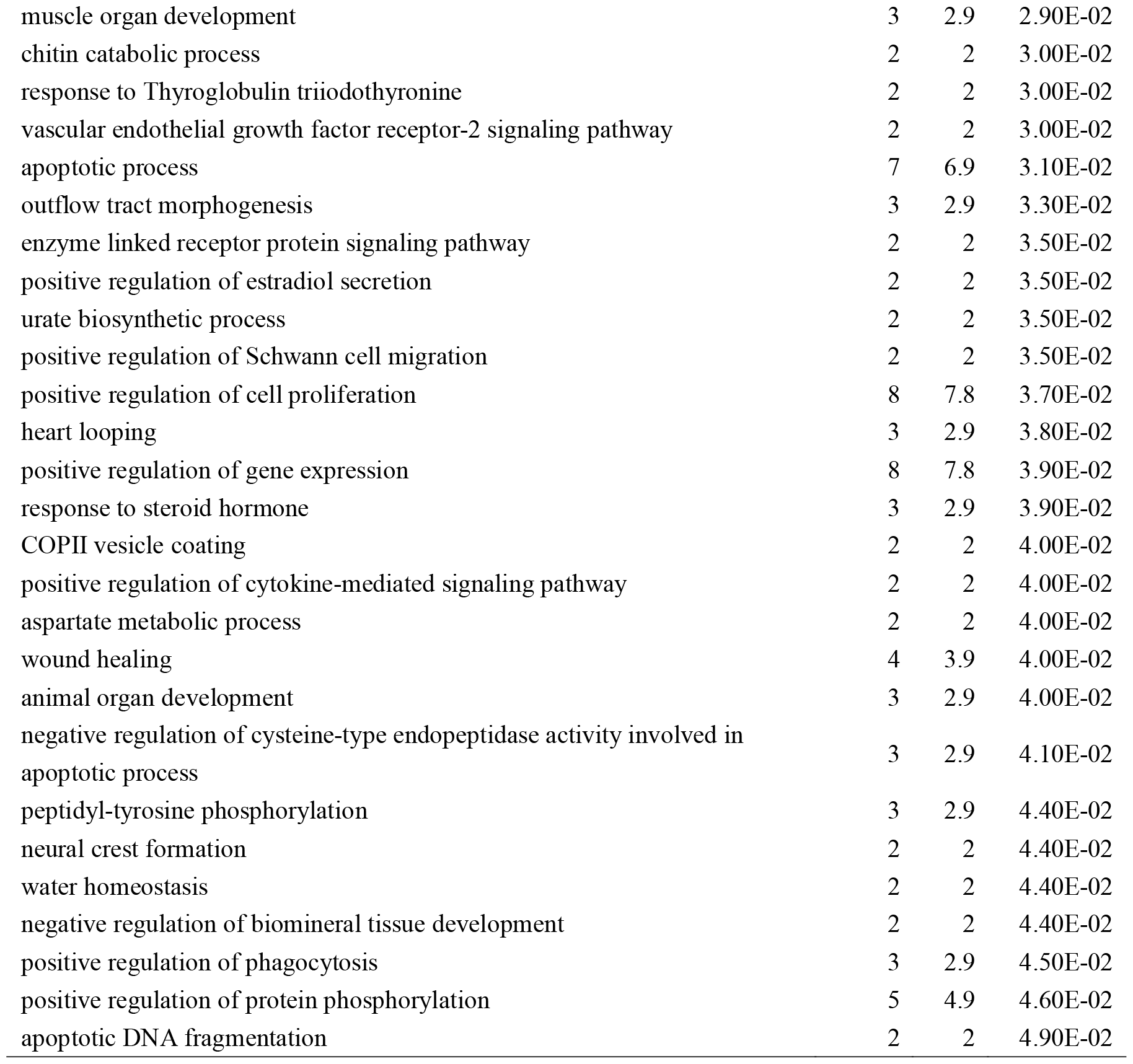
Biological processes (BP) to which differential proteins were enriched in the Zn-D0 and Zn-D4 groups (P-value < 0.05)

Zinc is essential for the proper functioning of the reproductive system. Differential proteins are enriched to biological processes including androgen catabolic processes, positive regulation of estradiol secretion, and response to steroid hormones. Zinc is essential for androgen expression^[64]^, estradiol affects zinc homeostasis in the follicle by controlling the expression of ZnT 9^[65]^.

Adequate intake of zinc reduces the risk of cardiovascular disease, and zinc plays a key role in maintaining vascular health by improving blood circulation and reducing arterial inflammation^[66]^. Differential proteins are enriched to biological processes including blood coagulation, heart development, aortic morphogenesis, cardiomyocyte proliferation, regulation of angiogenesis, vascular endothelial growth factor receptor-2 signaling pathway, and cardiac circulation.

Zinc is involved in neural signaling in the brain, aiding memory, learning, and cognitive functions^[1]^. Differential proteins are enriched to biological processes including acidification of synaptic vesicle lumen, positive regulation of neuroblast proliferation, cerebrospinal fluid secretion, regulation of Schwann cell migration, and neural crest formation.

Zinc is involved in the production and regulation of cells of the immune system and can help heal wounds^[67]^. Differential proteins are enriched to biological processes including the regulation of cytokine production involved in the inflammatory response, inflammatory responses, immune system processes, wound healing, and the regulation of phagocytosis.

Zinc protects cells from oxidative damage by free radicals and reduces oxidative stress^[1]^. Differential proteins are enriched to biological processes including positive regulation of reactive oxygen species metabolic processes and cellular response to hydrogen peroxide.

Zinc is important for regulating gene expression, DNA metabolism, chromatin structure, cell proliferation, maturation, and apoptosis^[68]^. Differential proteins are enriched to biological processes including regulation of the apoptotic process, cellular response to growth factor stimulation, regulation of cell death, apoptotic process, regulation of cell proliferation, apoptotic DNA fragmentation, regulation of cysteine-type endopeptidase activity involved in apoptotic process, and regulation of gene expression.

Zinc is a structural component of many proteins (e.g., many enzymes and transcription factors) and is involved in the metabolism of carbohydrates, proteins, and lipids, aiding in nutrient absorption^[1]^. Biological processes that differential protein enrichment to include response to organic matter, cellular chloride homeostasis, cellular sodium homeostasis, response to vitamin D, aspartate metabolic processes, and water homeostasis.

### 3.3 Molecular function and KEGG pathway analysis

Gene Oncology (GO) analysis of 112 differential proteins (P-value <0.05, FC>1.5 or <0.67) using the DAVID database enriched to 24 molecular functions (MF) (P-value <0.05), as shown in Table 3.

**Table 3.** Molecular functions (MF) enriched to differential proteins in Zn-D0 and Zn-D4 groups (P value < 0.05)

| Term | Count | % | P-Value |
| --- | --- | --- | --- |
| calcium ion binding | 18 | 17.6 | 1.60E-07 |
| integrin binding | 7 | 6.9 | 1.50E-04 |
| identical protein binding | 23 | 22.5 | 4.50E-04 |
| protease binding | 6 | 5.9 | 8.50E-04 |
| cuprous ion binding | 3 | 2.9 | 1.20E-03 |
| signaling receptor activity | 6 | 5.9 | 1.30E-03 |
| receptor binding | 8 | 7.8 | 4.70E-03 |
| cadherin binding | 4 | 3.9 | 5.70E-03 |
| proton-transporting ATPase activity, rotational mechanism | 3 | 2.9 | 1.00E-02 |
| protein binding | 18 | 17.6 | 1.00E-02 |
| deoxyribonuclease II activity | 2 | 2 | 1.10E-02 |
| folic acid receptor activity | 2 | 2 | 1.10E-02 |
| extracellular matrix binding | 3 | 2.9 | 1.30E-02 |
| methotrexate binding | 2 | 2 | 1.60E-02 |
| ATPase binding | 4 | 3.9 | 1.90E-02 |
| protein homodimerization activity | 10 | 9.8 | 1.90E-02 |
| chitinase activity | 2 | 2 | 2.10E-02 |
| cupric ion binding | 2 | 2 | 2.10E-02 |
| transmembrane receptor protein tyrosine kinase activity | 3 | 2.9 | 3.00E-02 |
| calcium-dependent phospholipid binding | 3 | 2.9 | 3.40E-02 |
| phosphatase activity | 3 | 2.9 | 3.90E-02 |
| thyroid hormone binding | 2 | 2 | 4.20E-02 |
| chitin binding | 2 | 2 | 4.20E-02 |
| copper ion binding | 3 | 2.9 | 4.70E-02 |

The molecular functions enriched to include cuprous ion binding, cupric ion binding, and copper ion binding. There are complex interactions between iron, copper and zinc^[70,69]^. Excessive zinc intake leads to copper deficiency, which leads to decreased iron absorption and ultimately anemia^[71]^.

Eighteen differential proteins were enriched for the molecular function of calcium ion binding, four differential proteins were enriched for the molecular function of calcineurin binding, and three differential proteins were enriched for the molecular function of calcium-dependent phospholipid binding. There appears to be an interaction signal between zinc and calcium ions^[5]^. Zinc homeostasis may be closely related to intracellular calcium signaling^[72]^.

Zinc ions are involved in extracellular signal recognition, signal transduction and second messenger metabolism^[67]^. The 6 differential proteins are enriched to the molecular function of signaling receptor activity. The molecular functions enriched to also include protease binding, receptor binding, ATPase activity, protein binding, and integrin binding.

Meanwhile, the biological processes that differential proteins are enriched to include response to calcium ions, positive regulation of ERK1 and ERK2 cascades, regulation of protein kinase B signaling, cellular phosphate ion homeostasis, regulation of kinase activity, regulation of protein tyrosine kinase activity, Notch signaling pathway, enzyme-linked receptor protein signaling pathway, peptidyl tyrosine phosphorylation, and positive regulation of protein phosphorylation.

Gene Oncology (GO) analysis of 112 differential proteins (P-value <0.05, FC>1.5 or <0.67) using the DAVID database enriched to 10 KEGG pathways (P-value <0.05), as shown in Table 4.

**Table 4.** KEGG pathways enriched to differential proteins in Zn-D0 and Zn-D4 groups (P value <0.05)

| Term | Count | % | P-Value |
| --- | --- | --- | --- |
| Complement and coagulation cascades | 5 | 4.9 | 3.40E-03 |
| Human papillomavirus infection | 9 | 8.8 | 3.50E-03 |
| Focal adhesion | 6 | 5.9 | 1.50E-02 |
| Collecting duct acid secretion | 3 | 2.9 | 1.60E-02 |
| ECM-receptor interaction | 4 | 3.9 | 2.50E-02 |
| Regulation of actin cytoskeleton | 6 | 5.9 | 2.60E-02 |
| Hematopoietic cell lineage | 4 | 3.9 | 2.70E-02 |
| Metabolic pathways | 19 | 18.6 | 3.20E-02 |
| PI3K-Akt signaling pathway | 7 | 6.9 | 3.90E-02 |
| Endocytosis | 6 | 5.9 | 4.50E-02 |

KEGG pathways enriched to include complement and coagulation cascades, Human papillomavirus (HPV) infection, ECM-receptor interactions, regulation of the actin cytoskeleton, hematopoietic cell profiles, metabolic pathways, and the PI3K-Akt signaling pathway.

Chronic zinc deficiency leads to a marked reduction in cell-mediated immunity, antibody response and antibody affinity, complement system and phagocytic activity^[73]^. High dietary zinc is negatively associated with high-risk HPV (hrHPV) infection^[74]^. In neuronal cells, zinc deficiency induces oxidative stress and alters the normal structure and dynamics of the cytoskeleton^[75]^. Zinc released from lysosomes is essential for ERK and PI3K / Akt activation^[67]^. Excessive zinc intake leads to copper deficiency, resulting in decreased iron absorption and ultimately anemia^[71]^.

## 4 Perspective

The results illustrate that short-term zinc gluconate supplementation affects the organism, and the urine proteome of rats can show changes in zinc-related proteins and biological functions, and also suggests that the urine proteome can comprehensively and systematically reflect the overall changes in the organism. The present study provides clues for an in-depth understanding of the metabolic process, mechanism of action, and biological function of zinc in organisms from the perspective of urine proteomics, as well as new research perspectives and methodological insights for future nutritional studies.

However, the supplement used in this study was zinc gluconate, but the correlation of differential proteins and biological functions with zinc was analyzed, not the effect of gluconate ions on the organism. On the one hand, the effect of zinc on the organism is more significant and easier to observe; on the other hand, there are fewer studies on the effect of gluconate ion on the organism. Due to limited resources, only one concentration of zinc gluconate was used in this study to gavage five rats, and subsequent studies may consider adding different supplement forms, adding different concentration gradients, increasing the number of samples, and expanding the study population. We expect that subsequent researchers will utilize the methodology and results of this study to make further additions to help translate and apply the experimental results for the benefit of human health.

